# Acute kidney injury disrupts cardiac remodeling via SerpinA3N

**DOI:** 10.1101/2025.09.19.677479

**Authors:** Gowda S. Pramod, Stanely Qu, Runze Ni, Nadezhda Zeleznova, Minghua Li, Nohely Hernandez, Lei Wang, Nathan Hall, Bi Zhao, Justin Gibbons, N Sanjib Banerjee, Ganesh V Halade, Lucian Lozonschi, Enyin Lai, Kristof Williams, Ruisheng Liu

**Affiliations:** Department of Molecular Pharmacology and Physiology, Morsani College of Medicine, University of South Florida (USF), Tampa, Florida, USA; Department of GenGHS, Genomics Program, College of Public Health, USF, Tampa, Florida, USA; Department of Biochemistry and Molecular Genetics, University of Alabama at Birmingham, Birmingham, AL 35294, USA; Division of Cardiovascular Sciences, Department of Medicine, Heart Institute, USF, 560 Channelside Dr, Tampa, FL 33602, USA; Division of Cardiothoracic and Transplantation Surgery, Tampa General Hospital (TGH), 1 Tampa General Circle, Tampa, FL 33606, USA; USF-TGH Transplant Research Center (UTRC), USF Health, USF, Tampa, Florida, USA

## Abstract

Cardiorenal syndrome type 3 (CRS-3) also known as acute reno-cardiac syndrome, refers to a condition where acute kidney injury (AKI) leads to acute cardiac dysfunction or injury. The underlying mechanisms have not been elucidated. In the current study, we examined the potential mechanisms for the development of myocardial pro-fibrotic factors and cardiac remodeling in C57BL/6J mice after AKI. The mice were subjected to bilateral pedicle clamping for 30 min followed by 24 h of reperfusion. Heart tissue was collected from control (Ctrl) and AKI mice and subjected to proteomic analysis by liquid chromatography-mass spectrometry (LC-MS) and differential mRNA expression analysis by RNA-seq. Cardiac tissue was collected to assess serine protease activity, RNA, protein and cytokine expression. The results indicate that AKI significantly upregulated SerpinA3N in the heart which was negatively correlated with serine protease (Granzyme B) activity. Further, AKI induced the development of several pro-fibrotic factors, induced mitochondrial dysfunction and increased inflammation. Finally, using H9C2 cells we demonstrated that inhibiting SerpinA3N with XAV939 increased Granzyme B activity. Thus, this study suggests that AKI leads to the development of pro-fibrotic factors in the heart and disrupts cardiac remodeling by SerpinA3N mediated inhibition of serine protease Granzyme B activity. Decreased SerpinA3N expression in the heart followed by AKI could lead to a more balanced extracellular matrix (ECM) composition in the heart with an alleviated pro-fibrotic response.

## Introduction

Acute kidney injury (AKI) is characterized by a rapid decline in kidney function over a short period, and is a common complication in hospitalized patients, with high morbidity and mortality (1-3). AKI can be caused by surgery, sepsis, trauma, nephrotoxic drugs, kidney stones, bladder cancer and enlarged prostate (4). Renal ischemia/reperfusion (I/R) is a critical factor that causes AKI, which often occurs after major surgeries such as kidney transplantation and cardiac surgery in multiple hospital settings (5,6). AKI frequently results in remote organ dysfunction involving the heart, lung, liver, intestines and brain through immune mediated inflammatory mechanisms (7-9).

Cardiorenal syndrome type 3 defines the syndrome in which AKI causes acute cardiac dysfunction (10). AKI can lead to both short- and long-term adverse effects on cardiac function (10-13) especially with existing cardiac complications such as heart failure, myocardial infarction (14,15) and post-cardiac surgery (16-18). It has previously been reported that AKI directly impairs cardiac function by reducing cardiac contractility in rodent models of AKI at 48 and 72 hours (19,20). Normal cardiac contractility requires ATP, and the preferred source is oxidative phosphorylation in mitochondria. Mitochondrial dysfunction is known to be associated with AKI-mediated heart injury (20), thus reduced ATP generation due to mitochondrial dysfunction is a fundamental mechanism of impaired cardiac contractility (21). Apart from mitochondrial dysfunction, AKI is known to trigger oxidative stress, apoptosis, inflammation and immune cell infiltration to the heart (22-24). Following cardiomyocyte necrosis and apoptosis, inflammatory cells recruited to the site of injury release cytokines that activate fibroblasts (25-28). Activated fibroblasts or myofibroblasts deposit extracellular matrix (ECM) components to form scar tissue at the injured regions in the heart (29). Cardiac fibrosis is the abnormal buildup of scar tissue in the myocardium, which can impair the heart’s ability to pump blood effectively, potentially leading to heart failure. The nascent scar tissue is susceptible to proteolytic degradation by a variety of proteases secreted primarily by infiltrating immune cells (30,31). Excessive proteolytic activity can lead to disruption of scar organization and impair cardiac remodeling (32), while reduced proteolytic activity can enhance scar tissue formation leading to fibrosis. The degree of fibrosis is determined by the balance between ECM deposition by myofibroblasts and the degradation of ECM by proteases secreted by invading immune cells.

Excessive protease activity in the injured region is regulated by a system of protease inhibitors. Serine protease inhibitors (Serpins) represent the largest and the most functionally diverse group of protease inhibitors and serve as critical regulator molecules in regulating protease activity in tissues (33,34). Emerging clinical studies have shown that SerpinA3 levels significantly increased after acute myocardial infarction and higher SerpinA3 levels were associated with major adverse cardiovascular events (35). However, little is known about the role of Serpins in regulating cardiac re-modelling after acute cardiac injury and after AKI. In this work, we demonstrate that, 24 hours after ischemic AKI, SerpinA3N is robustly expressed in the heart, while its target Granzyme B (GZMB) expression is also upregulated but with decreased serine protease activity in the heart. We also show that SerpinA3N directly prevents GZMB activity *in vitro* using H9C2 (Cardiomyoblast) cells. Further, we demonstrate that increased SerpinA3N expression is positively correlated with inhibition of Granzyme B activity and cardiac fibrosis molecular makers and thus might have a pivotal role in regulating post-AKI cardiac fibrosis.

## Materials and Methods

### AKI surgical procedure

Acute kidney injury (AKI) was induced by bilateral pedicle clamping in male C57BL/6J mice (average age: 12 weeks; weight: 25 g). Mice received a preoperative analgesic via subcutaneous injection 30 minutes prior to surgery; sustained-release Buprenorphine (1 mg/kg, SR, E72-hour formulation). Anesthesia was induced and maintained with 2–4% isoflurane in oxygen, and depth was verified by the absence of a pedal withdrawal reflex. All surgical procedures were conducted under aseptic conditions. A vertical midline laparotomy was performed to expose the abdominal cavity. The intestines were gently manipulated to the left and kept moist with sterile saline. The right kidney was exposed and using straight-tip ultra-fine forceps (Fine Science Tools, #11370-40), small openings were carefully created on both sides of the right renal pedicle to facilitate the later placement of a microvascular clamp (Fine Science Tools, #18055-04). The intestines were then repositioned to expose the left kidney, where identical preparation of the left renal pedicle was performed. Once both renal pedicles were prepared, bilateral renal ischemia was induced by placing microvascular clamps across each pedicle simultaneously. Ischemia was maintained for 30 minutes, during which body temperature was continuously monitored and maintained between 36.8°C and 37.2°C using a feedback-controlled heating pad. Following the ischemic period, the clamps were carefully removed to initiate reperfusion. The intestines were returned to the peritoneal cavity, and the abdominal wall was closed in two layers using 5-0 sutures: an absorbable polyglycolic acid suture for the muscular layer and a non-absorbable monofilament suture for the skin. Mice were allowed to recover on a heated surface and monitored until ambulatory. After a 24-hour recovery period, mice were euthanized, and tissues were collected for analysis. All experimental procedures were approved by the Institutional Animal Care and Use Committee at The University of South Florida (IACUC protocol #12005R) and performed in accordance with institutional and NIH guidelines for the care and use of laboratory animals.

### Glomerular filtration rate (GFR)

GFR was quantified in conscious *C57BL/6J* mice by transcutaneous clearance of fluorescein-isothiocyanate-sinistrin (FITC-sinistrin) using a dedicated fluorescence monitor (MediBeacon Preclinical MX Transdermal GFR Monitor; MediBeacon GmbH, Mannheim, Germany; Studio software v2.3). Measurements were obtained on day 0 (baseline, pre-ischemia) and day 1 (24 h post-ischemia-reperfusion AKI, immediately before euthanasia). For each session, mice were briefly anesthetized with 3 % isoflurane. Dorsal fur was removed with electric clippers followed by the application of depilatory cream (Veet, Reckitt Benckiser). The monitor was then placed over the clean, depilated area to start recording. After 5 minutes of baseline reading, FITC-sinistrin (7 mg 100 g⁻¹ body weight; 28 mg mL⁻¹ stock in sterile 0.9 % NaCl) was administered into the retro-orbital venous sinus (injection volume ≈ 0.25 µL g⁻¹). The mice were returned to their home cage for unrestricted movement during data acquisition until the system’s automated endpoint. FITC-sinistrin elimination curves were exported and fitted to the model within MediBeacon Studio, yielding GFR by bodyweight (mL min⁻¹ kg⁻¹).

### Plasma creatinine (Pcr)

For Pcr concentration measurements, tail-vein blood (20 µL) was collected into lithium-heparinized micro-hematocrit capillaries. Samples were kept on ice and centrifuged (12,000 × g, 4 °C, 10 min). 10 µL of plasma was transferred to 1.5 mL Eppendorf tubes, snap-frozen in liquid nitrogen, and stored at -80 °C freezer until analyzed. Plasma creatinine was quantified by isotope-dilution UHPLC-MS/MS at the UAB-UCSD O’Brien Center Biomedical Resource Core. (https://journals.physiology.org/doi/full/10.1152/ajprenal.00017.2020?rfr_dat=cr_pub++0pubmed&url_ver=Z39.88-2003&rfr_id=ori%3Arid%3Acrossref.org)

### Western blot analysis

Equal amounts of protein (50µg) were subjected to sodium dodecyl sulfate polyacrylamide gel electrophoresis (SDS-PAGE) using 4% to 12% gradient gels (Bio-Rad) and transferred to PVDF membranes (Thermoscientific Fisher). Transferred proteins were probed with appropriate antibodies (Table. 1) and detected using enhanced chemiluminescence reagents (Bio-Rad Clarity Western ECL Substrate). Band intensity was quantified by NIH ImageJ software version 1.54J.

### RNA isolation and real-time-qPCR

Total RNA was isolated using RNeasy Mini kits (Qiagen Inc.) according to the manufacturer’s specifications. cDNA was synthesized using SuperScript III First-Strand Synthesis System for RT-PCR (Invitrogen). Gene expression was determined by real-time RT-qPCR using Fast SYBR Green MasterMix (Applied Biosystems, Inc.) and gene-specific primers in an Quantstudio-Version 3 fast thermocycler. For each gene, expression levels were normalized to GAPDH. Experiments were performed in triplicate, and results are given as mean values SEM. Nucleotide sequences of primers are provided in Table 2.

**Table 1:**
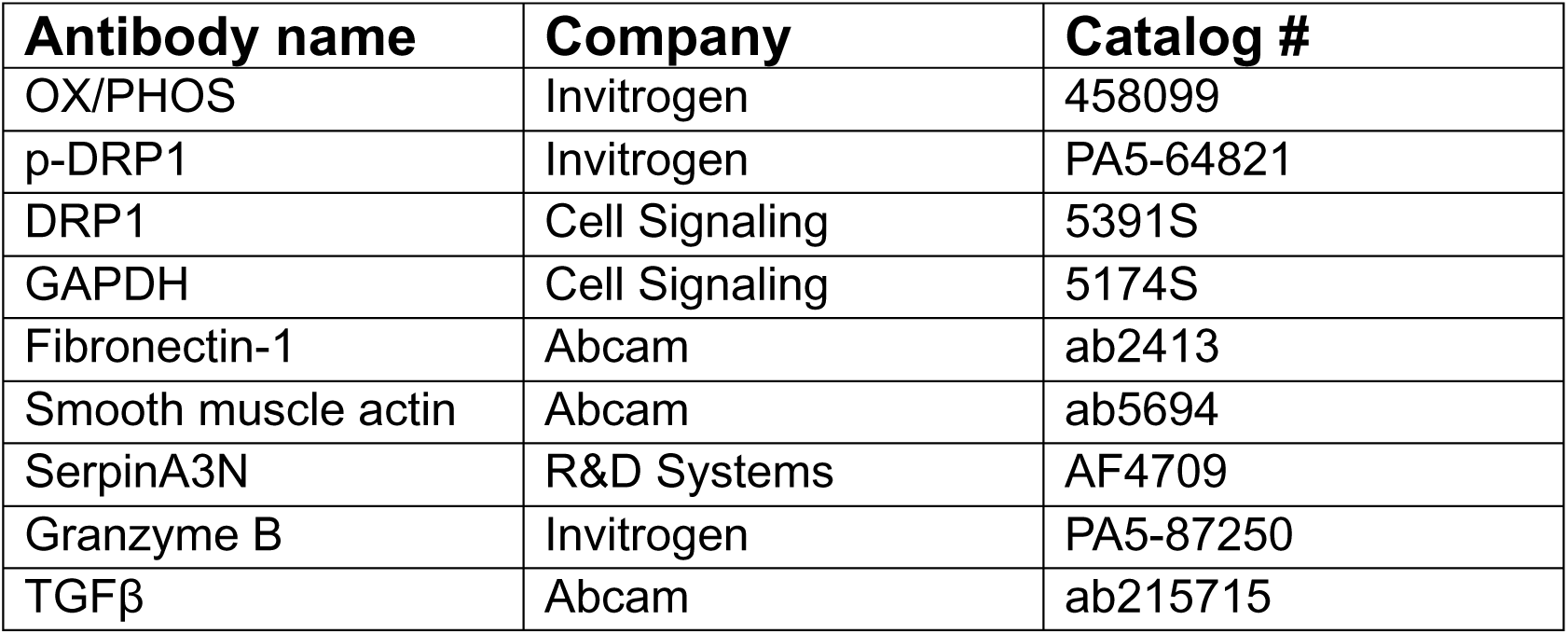
Antibodies list.

**Table 2:**
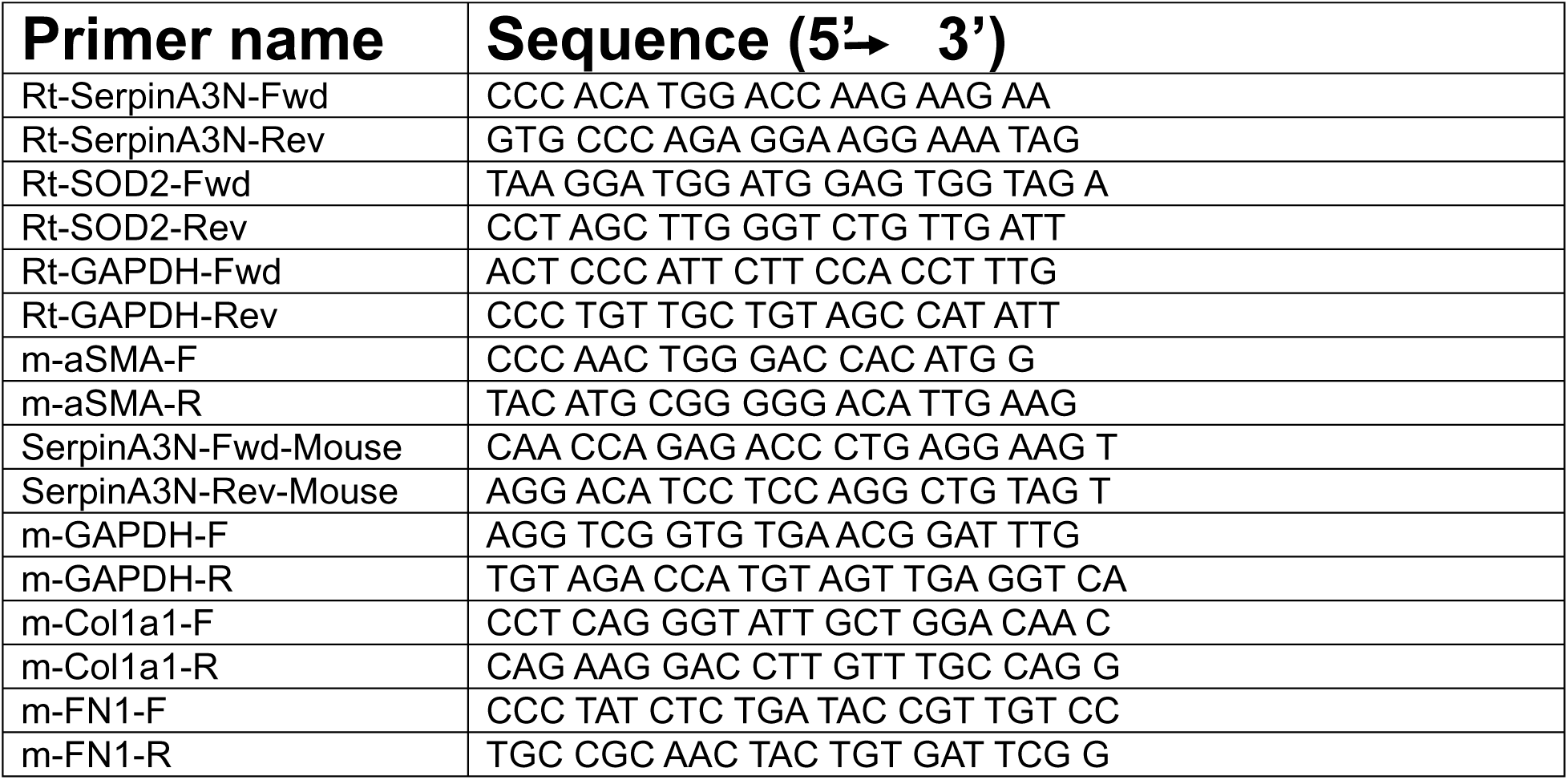
Real-time PCR Primers list.

### Cytokine Array Procedure

Cytokines were measured in frozen heart tissue collected 24 hours post 30-minute ischemia. Tissues were flash-frozen in liquid nitrogen and stored at -80°C until use. R&D Systems’ Proteome Profiler Array / Mouse XL Cytokine Array Kit (Catalog No. ARY028) was used. The manufacturer’s provided protocol was followed precisely. In brief, tissues were homogenized in PBS + 1% protease inhibitors using TissueLyser LT bead homogenizer (two 1-minute cycles at 50 oscillations per second). Samples were centrifuged (10,000 x g for 5 minutes), supernatant was collected, and protein concentration was measured by Bio-Rad Protein Assay (5000006, Bio-Rad). Protein concentrations were diluted to 200ug for final samples. Regarding membrane preparation, membranes were first blocked. 2 mL of the provided blocking buffer (buffer 6) was added to the membranes, followed by incubation at room temperature for 1 hour on a rocking platform shaker. Buffer 6 was aspirated from the wells, and the samples diluted in array buffer (buffer 4) were added to the membranes and incubated overnight at 2-8°C on a rocking platform shaker. Membranes were washed with the provided wash buffer (3 x 10-minute washes per wash cycle). The antibody detection cocktail was added to the membranes and incubated at room temperature for 1 hour on a rocking platform shaker. Membranes were washed, Streptavidin-HRP was added to the membranes, and incubated at room temperature for 30 minutes on a rocking platform shaker. Membranes were washed, and the provided Chemi-Reagent Mix was applied to the membranes and incubated for 1 minute immediately before imaging. Upon assay completion, the membranes were imaged with the Bio-Rad ChemiDoc MP imaging system. Membrane analysis, quantification, and densitometry of positive markers were completed using Bio-Rad Image Lab 6.1 software.

### ELISA

Whole blood from Ctrl and AKI mice was collected in an EDTA coated tube and spun at 12000g for 10 minutes at 4deg, and the supernatant containing plasma was used for ELISA. The remaining steps were the same as those for the detection of BNP (RayBiotech, United States, Cat# EIAM-BNP) and Mouse Cardiac Troponin T (Antibodies.com LLC, United States, Cat# A73988) followed according to the manufacturer’s protocol.

### Histological Staining & Imaging

Samples were fixed in 4% paraformaldehyde (PFA) for 24 hours, followed by immersion in 70% ethanol for an additional 24 hours. Paraffin embedding, slide preparation, and hematoxylin and eosin (H&E) staining were performed by the Small Animal Imaging Laboratory Core at the H. Lee Moffitt Cancer Center & Research Institute. Whole tissue section images were acquired at magnifications ranging from 4× to 20× using the Olympus VS120 slide scanning system (Olympus, Tokyo, Japan), provided by the Lisa Muma Weitz Imaging Core Facility at the University of South Florida (Tampa, FL).

### Proteomics digest

Tissue samples were lysed in 5% SDS/ 50mM TEAB buffer with 1% Halt protease inhibitor cocktail, after centrifugation supernatant were transferred to a sperate tube and a Peirce 660nm Protein plus assay performed to determine protein concentration. 25 ug of each sample was digested using a S-TRAP micro column from Protifi (#C02-micro-80) according to the supplied SOP. Digested samples were dried and then rehydrated in 1% Formic acid for Lc-MS analysis (36).

### LC-MS analysis

Peptides were characterized using a Thermo Q-exactive-HF-X mass spectrometer coupled to a Thermo Easy nLC 1200. Samples separated at 300nl/min on an Acclaim PEPMAP 100 trap (75uM, 2CM, c18 3um, 100A) and a Thermo easy spray column (75um, 25cm, c18, 100A) using a 120-minute gradient with an initial starting condition of 2% B buffer (0.1% formic acid in 90% Acetonitrile) and 98% A buffer (0.1% formic acid in water). Buffer B increased to 28% over 100 minutes, then up to 40% in an additional 10 minutes. High B (90%) was run for 15 minutes afterwards. The mass spectrometer was outfitted with a Thermo nano spray easy source with the following parameters: Spray voltage: 1.85V, Capillary temperature: 275dC, Funnel RF level=40. Parameters for data acquisition were as follows: for MS data the resolution was 60,000 with an AGC target of 3e6 and a max IT time of 50 ms, the range was set to 400-1600 m/z. MS/MS data was acquired with a resolution of 15,000, an AGC of 1e4, max IT of 50 ms, and the top 30 peaks were picked with an isolation window of 1.6m/z with a dynamic execution of 25s. For data analysis, resulting samples were processed using Max quant 2.0.3.1 free software (37). Reviewed Human database was downloaded from Uniprot and searched with the following parameters: tryptic enzyme with a max of 2 missed cleavages, a precursor mass tolerance of 10ppm and a fragment mass tolerance of 0.02 Da.

Modifications included Oxidation, Acetyl, and Carbamidomethyl and pSTY. FDR rate was set at 0.01, LFQ intensities were compared between samples for ID. Perseus analysis software was used to sort and filter data (38). Identifications were filtered to be found in at least 2/3 samples in at least 1 group.

### Transcriptome RNA sequencing

Total RNA from Ctrl and AKI heart tissues was isolated using the RNeasy Mini Kit (Qiagen, Germany). After assessment of the integrity of total RNA using the Agilent 2100 Bioanalyzer (Agilent Technologies Inc., USA), the samples with RNA integrity number > 7.0 were used for sequencing. RNA was sent out to GENEWIZ from Azenta Life Sciences (South Plainfield, NJ, USA) for RNA sequencing (RNA-seq) experiment. Differential expression analysis of mRNA was performed using the R package edgeR.

### H9c2 cell culture and *in vitro* oxidative stress induction using H_2_O_2_

H9c2 cells were originally derived from embryonic rat ventricular cardiomyocytes and were a kind gift from Dr. Jin O’Chi at University of South Florida (USF). Cardiomyocytes were cultured in Dulbecco’s modified Eagle’s medium (GIBCO, MA, USA) supplemented with 10% fetal bovine serum (Biological Industries, CT, USA) and 5% penicillin and streptomycin (GIBCO, MA, USA). The cultures were maintained in a humid incubator (95% air and 5% CO_2_) at 37 °C. To induce oxidative stress, the cells were treated with H_2_O_2_ at a final concentration of 20µM for 24hrs.

### Granzyme B activity assay

We used Abcam’s Granzyme B Inhibitor Screening Assay Kit (Cat# ab157405), heart tissue lysates from Ctrl and AKI were used in the experiment as a source of active Granzyme B (GZMB). Active GZMB hydrolyzes the GZMB substrate provided in the kit to release the quench of fluorescent group, which can be detected fluorometrically at Ex/Em = 380/500 nm. The RFU of fluorescence generated by hydrolyzation of substrate is directly proportional to the activity of GZMB.

### IPA

From the IPA website: (https://digitalinsights.qiagen.com/citation-guidelines/)

Data were analyzed with the use of QIAGEN IPA Interpret (QIAGEN Inc., https://digitalinsights.qiagen.com/IPA).

The [networks, functional analyses, etc.] were generated through the use of QIAGEN IPA Interpret (QIAGEN Inc., https://digitalinsights.qiagen.com/products-overview/discovery-insights-portfolio/analysis-and-visualization/qiagen-ipa/qiagen-ipa-interpret/) (1D).

To investigate the biology underlying the molecular function and disease association, Ingenuity Pathway Analysis (IPA) (Qiagen, Redwood City, CA) software was used in the analysis. IPA analysis identified canonical pathways in which differentially expressed features were significantly over-represented, based on a hypergeometric/right-tailed Fisher’s exact test with false discovery rate (FDR)-adjusted p-value <0.05 (39).

### Statistical analysis

All data were presented as mean ± SEM of at least three independent experiments. All analyses were performed with GraphPad Prism 7 software (GraphPad, USA). The normality assumption of the data distribution was assessed using the Shapiro-Wilk test. Unpaired two-tailed Student’s *t* tests were used to determine the statistical difference between two groups. One-way analysis of variance (ANOVA) followed by Turkey’s post hoc test was performed to determine the statistical difference between multiple groups with one variable.

## Results

### IRI surgery causes significant kidney injury

Male mice were assigned to both Ctrl and AKI groups. In the case of Ctrl, laparotomy was performed to expose the abdominal cavity followed by closing the abdominal walls with an absorbable polyglycolic acid suture for the muscular layer and a non-absorbable monofilament suture for the skin. Bilateral acute kidney injury was induced in AKI mice, their pedicles were clamped for 30min to cause ischemia and following the ischemic period, the clamps were carefully removed to initiate reperfusion, and mice were allowed a 24-hour recovery period. All surgical procedures were conducted under aseptic conditions and under anesthesia. After a 24-hour recovery period, mice were euthanized, and blood and tissues were collected for analysis (Fig. 1A). Mice from both Ctrl and AKI groups were subjected to glomerular filtration rate (GFR) quantification at 0 and 24hr time points, AKI mice showed a significant decrease in GFR at 24hr after AKI surgery as compared to Ctrl control group (Fig. 1B). The upregulation of plasma creatinine (PCr) (Fig. 1C) as well as damaged renal tubules (Fig. 1D) was also observed in AKI mice as compared to Ctrl control group. These data indicate the development of AKI with significant kidney injury.

**Figure 1.**
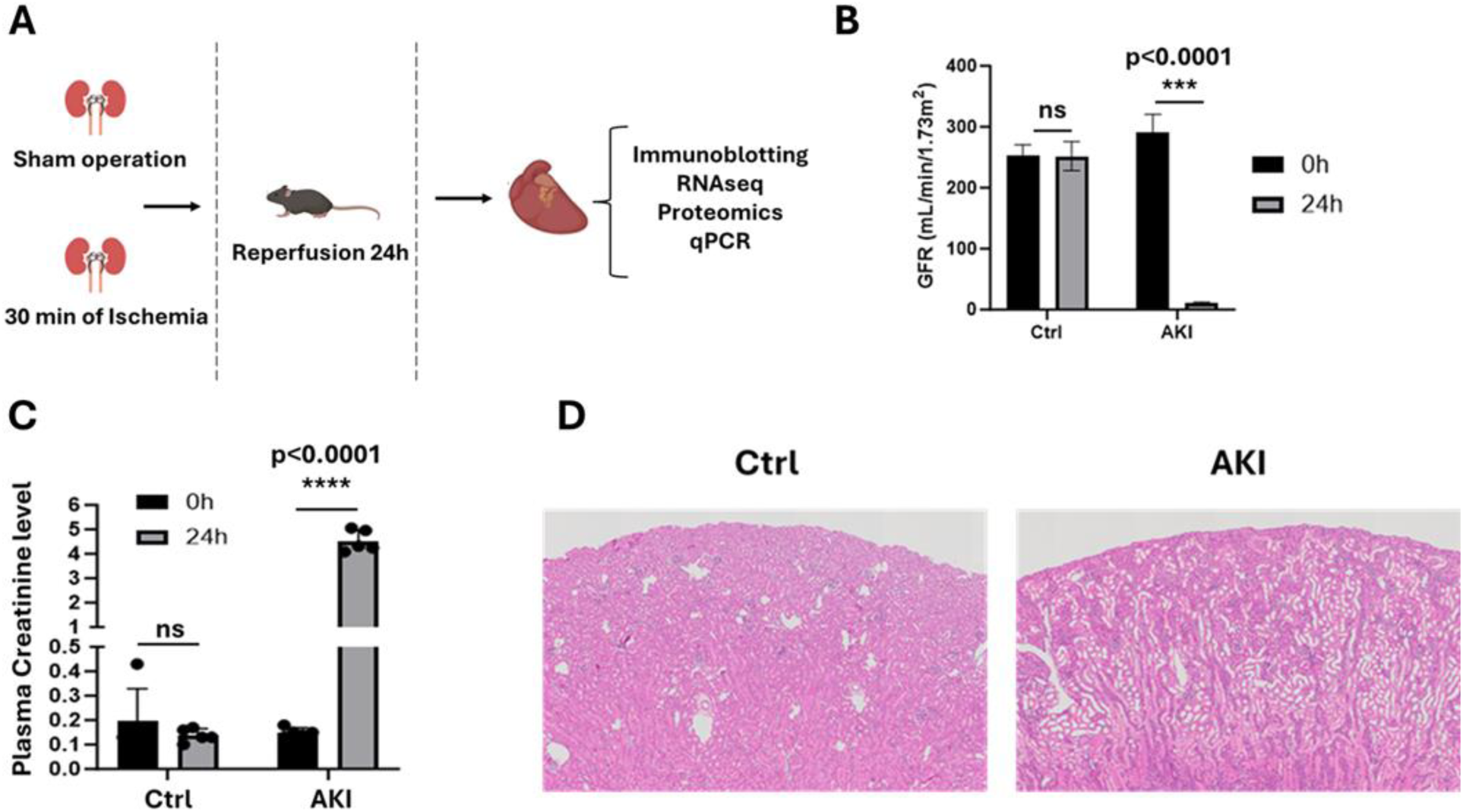
AKI surgery induces significant renal injury. A) Experimental design, C57BL/6J wild-type mice underwent a laparotomy and clamping of the renal artery and vein for 30 min, followed by 24 h of reperfusion. Blood, Kidney and Heart issues were collected for ELISA, immunoblotting, RNA, proteomics and RNA-seq analysis. B) GFR measurement was performed on the mice at Day 0 and Day 1 after AKI surgery. C) Plasma Creatinine levels were measured similarly at Day 0 and Day 1 after AKI surgery. D) Kidney tissues were histologically H&E stained to detect tubular damage after AKI. The P values were obtained with two-tailed Student *t* test (\**P* < 0.05; \*\**P* < 0.01; \*\*\**P* < 0.0001).

### CRS-3 is featured by Cardiac Dysfunction

The mouse CRS-3 model was established (Fig. 1A), and to evaluate cardiac damage, myocardial injury markers such as blood Troponin T and BNP were determined through ELISA. Following AKI, the levels of Troponin T and BNP were significantly increased when compared to that in the Ctrl group (Figs. 2 A, B). Besides that, cardiac inflammation was increased as assessed by RNA-seq and cytokine array analysis, pro-inflammatory cytokines such as IL-1b, IL-6, CCL2, IL-7, CSF-3 and ICAM-1 were increased and anti-inflammatory cytokines such as IL-4 and IL-17d were decreased in the heart tissues of AKI group as compared to the heart tissues of Ctrl group (Figs. 2 C, D). Given the indispensable role of mitochondria in regulating myocardial ATP supply, we questioned whether AKI is followed by a disruption of mitochondrial function. To achieve this aim, cardiac tissue from both Ctrl and AKI groups were subjected to RNA-seq analysis, which showed a significant decreased expression in mitochondrial electron transport chain (ETC) complexes (I, II, III, IV and V), which are crucial for cellular energy production through oxidative phosphorylation (Fig 2. E). ETC complex downregulation was further confirmed by western blotting using OX/PHOS antibody (Fig 2. F). Finally, we looked at mitochondrial division by analyzing p-DRP1 levels between Ctrl and AKI groups. It has been demonstrated that the increased p-DRP1 represents increased mitochondrial fission with less ATP production (40), our results confirmed that AKI groups showed an elevated p-DRP1 levels as compared to Ctrl (Fig 2. G). Taken together, our results confirmed that cardiac damage following AKI has been established by an increase in inflammation and mitochondrial dysfunction along with the classical cardiac injury markers TnT and BNP upregulation.

**Figure 2.**
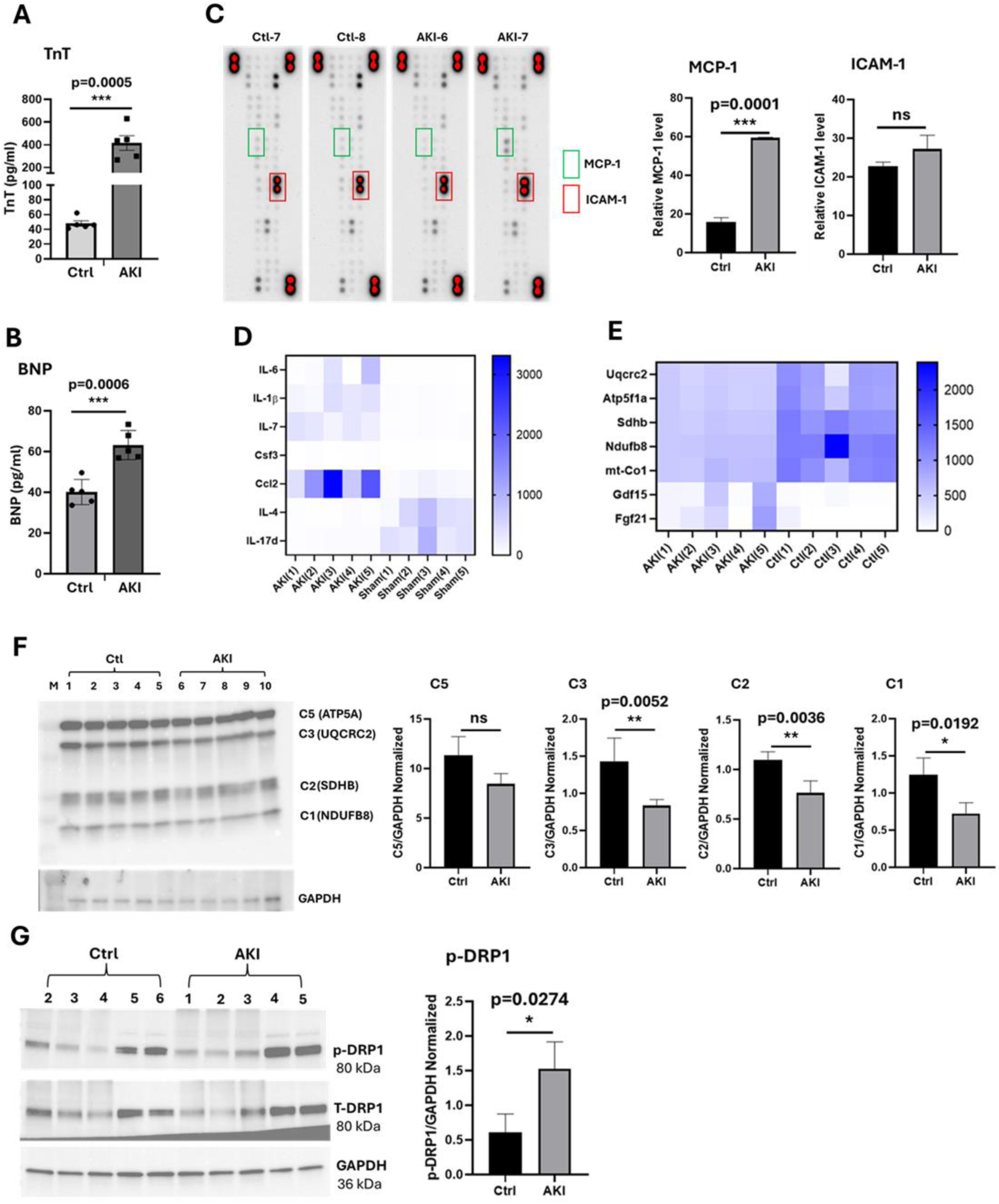
Renal AKI induces cardiac damage. A, B) Levels of Serum TnT and BNP from mice after AKI assessed by ELISA. C, D) Cytokine array and RNA-seq analysis of pro-inflammatory and anti-inflammatory gene expression between Ctrl and AKI groups. E, F) RNA-seq and Western blot analysis of mitochondrial ETC complexes (I, II, III, IV, V) between Ctrl and AKI groups. G) Western blots were used to analyze the expression of phosphorylated Drp1 (p-Drp1). The relative expression of p-Drp1 was evaluated in Ctrl and AKI. The P values were obtained with two-tailed Student *t* test (\**P* < 0.05; \*\**P* < 0.01; \*\*\**P* < 0.0001).

### Renal AKI induces pro-fibrotic factors in the heart

To determine the expression of pro-fibrotic factors after renal AKI, we analyzed transcriptomic data sets of hearts of adult male mice (C57Bl/6) subjected to acute AKI and observed increased expression of FN1, Col1a1, TGFb1, Acta2, Has1, Has2, TIMP1 and MMP3, while decreased expression of MMP13 in the injured heart (compared with the uninjured heart) 24hrs post-injury (Fig. 3A). We validated the RNA-seq data by Western blotting for FN1, TGFb1 and Acta2 (Fig. 3B), by qPCR for Col1a1, Acta2 and FN1 (Fig. 3C) and by cytokine array for TIMP1 (Fig. 3D)., Our IPA analysis revealed an enrichment of cardiac fibrosis genes in AKI as compared to Ctrl heart tissues (Fig. 3E). These findings conclusively demonstrate that renal AKI leads to an increase in pro-fibrotic factors in the heart.

**Figure 3.**
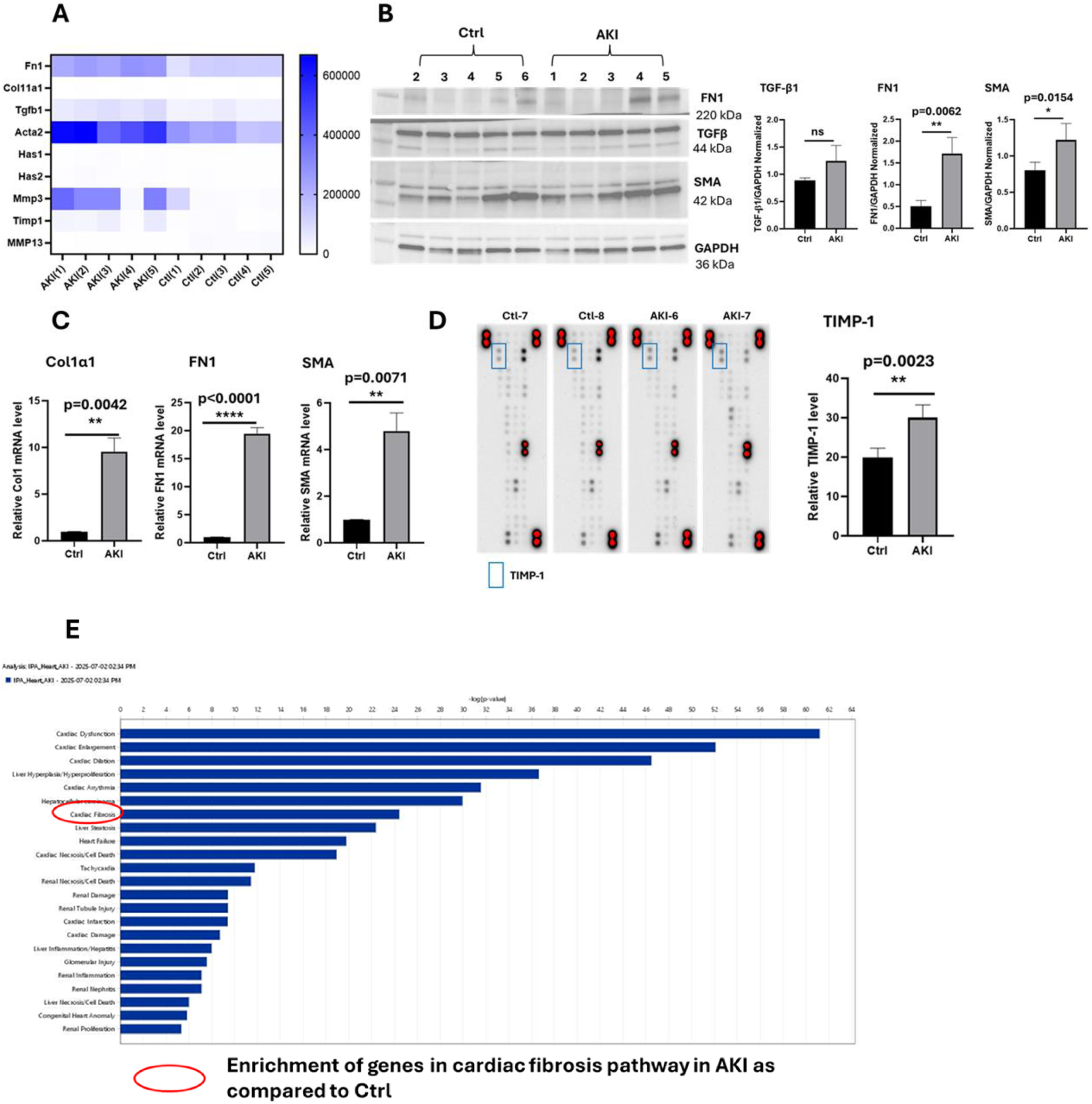
Renal AKI induces pro-fibrotic factors in the heart. A) RNA-seq analysis between Ctrl and AKI groups revealed an increase in pro-fibrotic gene expression, whereas MMP-13 anti-fibrotic gene was decreased in AKI. B) Western blot analysis of pro-fibrotic genes TGF-β1, FN1 and SMA between Ctrl and AKI groups. C) qPCR analysis of pro-fibrotic genes Col1A1, FN1 and SMA between Ctrl and AKI groups. D) Cytokine array analysis revealed increased TIMP-1 expression in AKI heart tissue as compared to Ctrl. E) Bar graph of Differentially regulated pathways obtained from IPA analysis. AKI group shows an enhanced cardiac fibrosis gene pathway alterations as compared to Ctrl. The P values were obtained with two-tailed Student *t* test (\**P* < 0.05; \*\**P* < 0.01; \*\*\**P* < 0.0001).

### Myocardial SerpinA3N Expression is Increased Following AKI

To better clarify the molecular mechanism that is activated by AKI and participates in acute myocardial damage, we performed LC-MS to understand without bias protein expression in the heart following AKI and in control. The results demonstrated that a total of 44 proteins (p < 0.05, log2 fold change) were upregulated or downregulated in heart tissue following AKI (Figures 4 A, B). Previous study had demonstrated that SerpinA3N, a serine-protease inhibitor is up-regulated in the context of myocardial infarction (MI) and is associated with regulating scar formation after MI (41). Our LC-MS results also showed a significant increase in SerpinA3N (Fig. 4C), we further validated with Western blotting and q-PCR for SerpinA3N expression in the heart tissues of AKI and Ctrl groups (Figs. 4D, E). A molecular function description of Gene Ontology (GO) further illustrated that 6 of the 44 proteins were involved in protease inhibitor activities (such as serine-type endopeptidase inhibitor activity, peptidase inhibitor activity, endopeptidase inhibitor activity, endopeptidase and peptidase regulator activity) (Figure 4F). With the help of *in silico* STRING analysis of protein–protein interactions, we found that SerpinA3N is involved in the crosstalk and signal integration of various molecular events in the heart after AKI (Figure 4G). Molecular function description of GO predicted that elevated SerpinA3N was correlated to inhibition of serine proteases in the heart of AKI as compared to Ctrl. Taken together, our data demonstrates an increased and active SerpinA3N function in the heart of AKI could inhibit serine proteases leading to increased scar tissue formation, disruptive cardiac remodeling and cardiac fibrosis.

**Figure 4.**
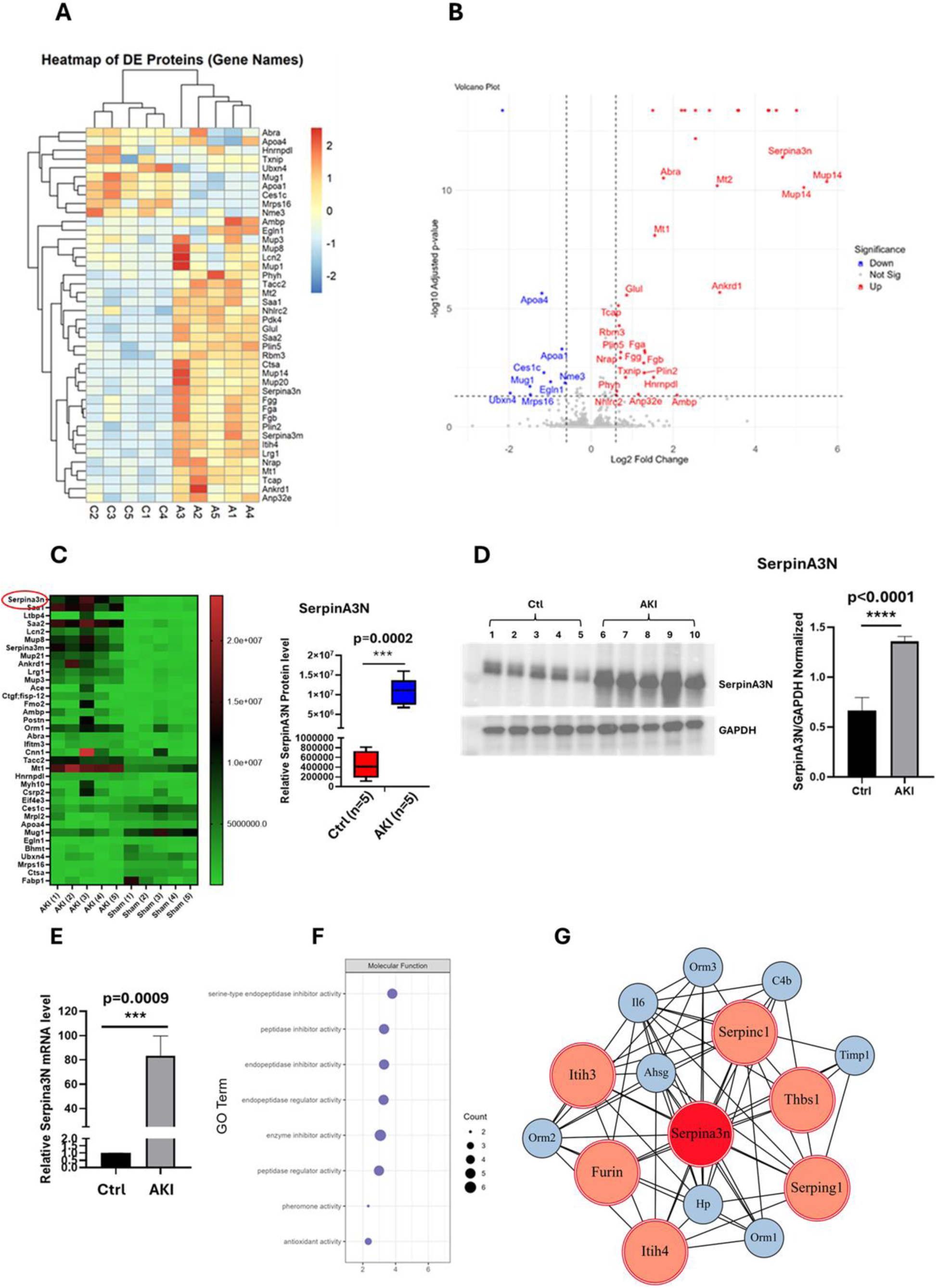
Renal AKI upregulates serine protease inhibitor protein SerpinA3N expression and activity in heart tissues. A, B) We processed proteomic data using *DEP* (52) version 1.28.0 and identified differentially expressed proteins between two conditions using function *test_diff* from *DEP* package. The results demonstrated that 44 proteins (adj.p < 0.05, absolute log2 fold change >=0.6) were upregulated or downregulated in heart tissues. C) SerpinA3N protein expression was significantly upregulated in the heart of AKI tissues as compared to Ctrl as identified by LC-MS. D, E) Western blot and qPCR analysis of SerpinA3N expression between Ctrl and AKI groups. F) Gene Ontology annotation was used to identify the molecular function of differentially expressed proteins. G) The protein–protein interaction network among the differentially expressed genes in heart tissue, the interaction was collected from STRING with median confidence 0.4. The P values were obtained with two-tailed Student *t* test (\**P* < 0.05; \*\**P* < 0.01; \*\*\**P* < 0.0001).

### SerpinA3N inhibits Granzyme B (GZMB) activity in the heart from AKI mice model

SerpinA3N is known to inhibit several serine proteases including granzyme B which is released by a variety of cytotoxic and non-cytotoxic cells (42). Granzyme B is known to be elevated after acute cardiac and neurological injury and contributes to matrix breakdown and ECM remodeling (43). The expression of GZMB was examined by Western blotting and the results demonstrated that AKI increased the expression of GZMB in the heart tissues as compared to Ctrl (Fig. 5A). However, GZMB activity significantly decreased in the AKI heart tissues as compared to Ctrl (Fig. 5B). Further, we observed an inverse correlation between SerpinA3N expression levels and GZMB activity, with AKI condition showing the highest SerpinA3N expression with significantly reduced GZMB activity as compared to Ctrl (Fig. 5C). Taken together, our results demonstrate that, though GZMB expression is increased in the heart of AKI group compared to Ctrl, their activity is significantly decreased, and this is due to the significantly high expression of SerpinA3N in the heart of AKI group in comparison to Ctrl.

**Figure 5.**
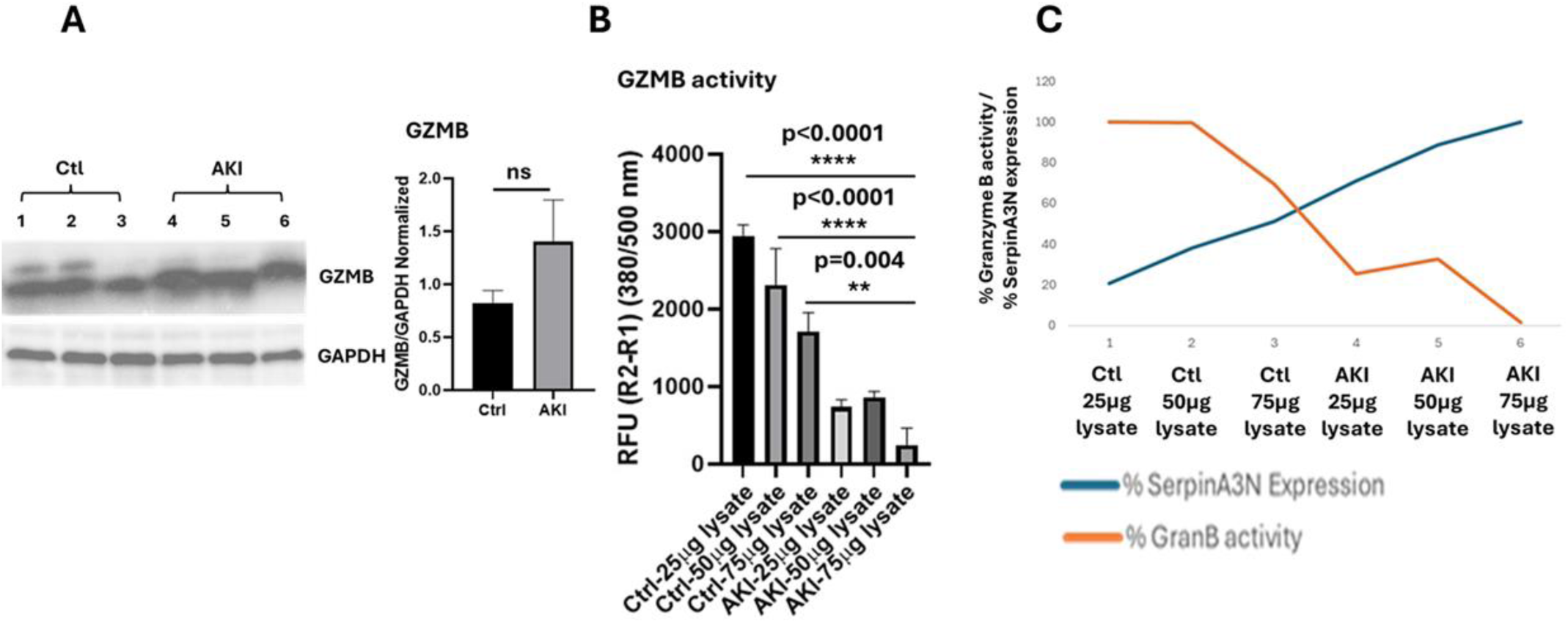
SerpinA3N expression in inversely correlated to Granzyme B (GZMB) activity. A) Western blot assessment of GZMB expression in the hearts of Ctrl and AKI mice. B) Increasing concentrations of Ctrl and AKI heart lysates were used to assess the GZMB activity, lysates were the source of SerpinA3N and GZMB proteins, the total lysate was treated with GZMB peptide substrate, which is hydrolyzed by GZMB, this hydrolysis rate was used to quantify GZMB activity. C) GZMB activity was plotted against SerpinA3N protein concentration, higher the SerpinA3N concentration the lower was GZMB activity and this negative correlation was higher in the AKI as compared to Ctrl. The P values were obtained for n=3 with an ANOVA and Tukey post-test used for experimental comparisons (\**P* < 0.05; \*\**P* < 0.01; \*\*\**P* < 0.0001).

### SerpinA3N directly inhibits GZMB activity *in vitro*

To assess if GZMB activity is directly inhibited by SerpinA3N and this is not an indirect inhibition. We employed H9c2 (Cardiomyoblast) cells, where these cells were stressed with H_2_O_2_ to induce oxidative stress (assessed by SOD-2 upregulation) and oxidative stress upregulates SerpinA3N expression in H9c2 cells (Fig. 6A), and SerpinA3N get secreted into the media. To understand if SerpinA3N directly inhibits GZMB activity, we blocked SerpinA3N function using XAV939 compound (Selleckchem, Cat # S1180) a known inhibitor of Wnt signaling (44). Transcription factor, C/EBP, increases Serpin transcription (45), while elevated Wnt signaling increases C/EBP expression and Serpin expression (46). A small molecule XAV939 inhibits Wnt signaling and negatively affects SerpinA3N function (44, 47). Since there are no direct inhibitors of SerpinA3N we used XAV939 to regulate SerpinA3N expression and function via inhibiting Wnt signaling with XAV939 compound. H9c2 cells 5X10^5^ were treated with H_2_O_2_ and XAV939 for 24hrs at various concentrations. H9c2 cells with no treatments were used as controls. Cell conditioned media (CM) from H9c2 cells from both treated and untreated groups were concentrated to one fifth of its original volume using AmiconYM-10 Centricon filters (molecular mass cutoff of 10 kDa; Millipore) for 90 min at 7000 rpm (4°C) down to a volume of 600ul (6X concentration). The concentrated CM was stored at 4°C until use. Concentrated CM was used as the source of SerpinA3N in the *in vitro* assay to assess GZMB activity. CM from cells treated with both H_2_O_2_ and XAV939 showed an increase in GZMB activity as compared to CM from cells treated with only H_2_O_2_ (Fig. 6B). These data indicate that XAV939 mediated inhibition of SerpinA3N has less inhibitory effect on GZMB activity. Thus, GZMB activity is directly affected by SerpinA3N expression, as it is known that GZMB physically interacts with SerpinA3N and their interaction can proceed along either the inhibitory or substrate pathways (48).

**Figure 6.**
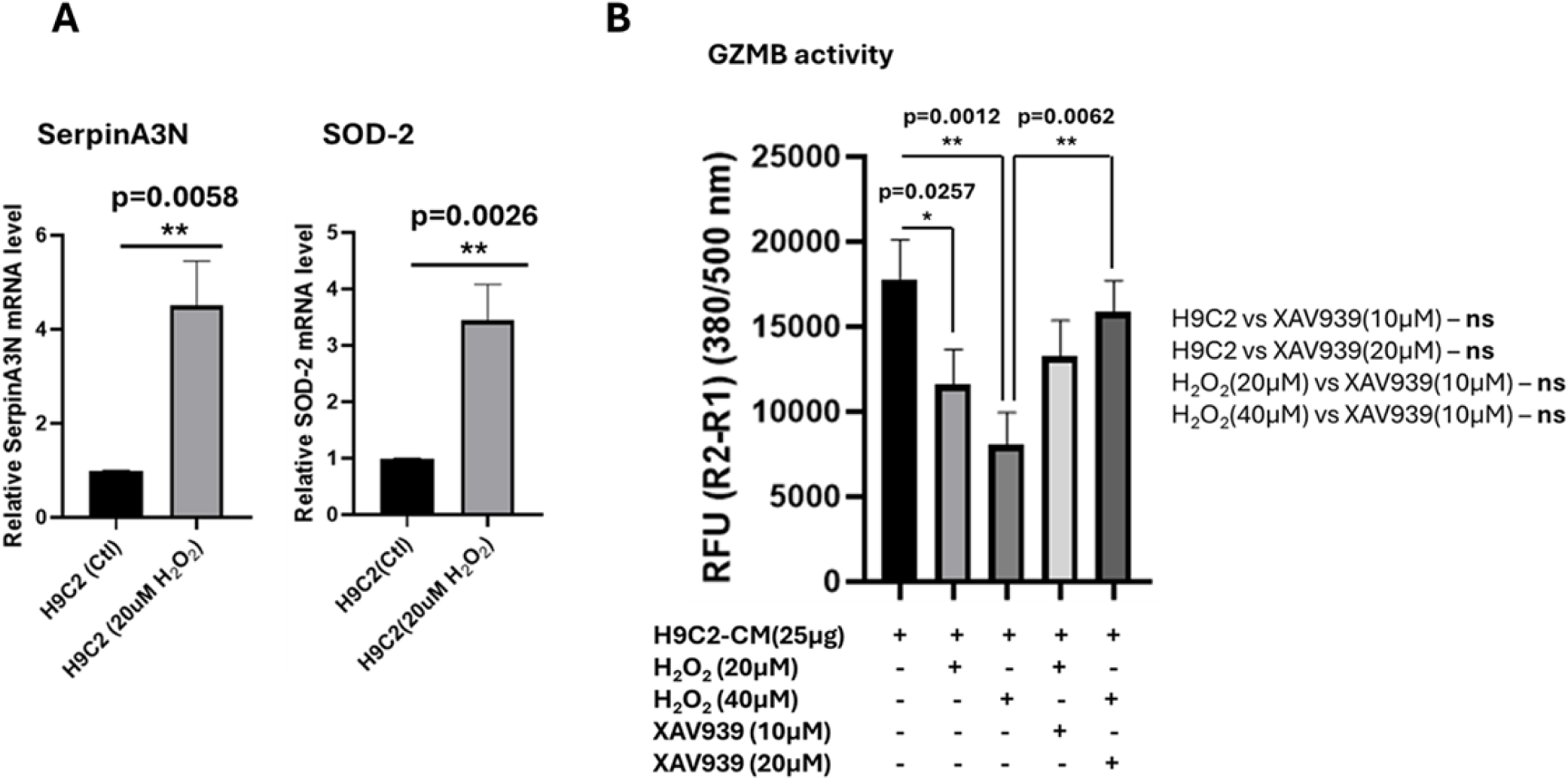
Inhibiting SerpinA3N increases Granzyme B activity. A) H9c2 cells were treated with H_2_O_2_ and assessed for oxidative stress marker SOD-2 and SerpinA3N expression by qPCR. B) H9c2 cell-conditioned media (CM) was concentrated after H9c2 cells were treated with H_2_O_2_ and XAV939 and the respective CM was assessed for GZMB activity with the GZMB inhibitor screening assay kit (ab157405) as described before. The P values were obtained for n=3 with an ANOVA and Tukey post-test used for experimental comparisons (\**P* < 0.05; \*\**P* < 0.01; \*\*\**P* < 0.0001).

## Discussion

The major findings of this study are that kidney injury results in the overexpression of serine protease inhibitor SerpinA3N and serine protease Granzyme B (GZMB) in the heart, however GZMB activity is significantly decreased by SerpinA3N in the heart. Kidney injury also results in increased pro-fibrotic factors in the heart.

To understand the relationship between SerpinA3N and pro-fibrotic factors, we used the IPA molecular activity predictor (MAP) analysis of the principal SerpinA3 interaction network (Fig. S1). The principal interaction network of SerpinA3 predicted that SerpinA3 indirectly dephosphorylates GSK3α and GSK3β and leads to their activities that contribute to an increase in fibrosis in the heart (49,50). Our RNAseq analysis showed an increase in GSK3α and GSK3β in the hearts of AKI as compared to Ctrl, suggesting a role for SerpinA3N in activating GSK3α and GSK3β and leading to pro-fibrotic condition, however our proteomic analysis did not detect the phosphorylation status of these proteins (data not shown), further investigation into these proteins is necessary to understand their interaction with SerpinA3N leading to pro-fibrotic condition in the heart.

Importantly, our study demonstrated that AKI increased inflammation and TGF-β1 expression in the heart which correlated to a significant increase in SerpinA3N expression and pro-fibrotic gene expression (Figs. 3, 4). This finding is consistent with a recent study (51) which showed that TGF-β1 induced SerpinA3N expression and pro-fibrotic genes expression in hiPSC-derived cardiac fibroblasts (Fig. S2). Previous studies from other laboratories and our current study, suggest that AKI-induced TGF-β1 increases SerpinA3N and pro-fibrotic factors in the heart and this could negatively affect cardiac remodeling leading to cardiac injury after AKI.

In this study, we established an AKI mouse model by subjecting the mice to bilateral pedicle clamping for 30 min followed by 24 h of reperfusion (Fig. 1) and used this model to demonstrate kidney and cardiac injury and understand the underlying mechanisms of cardiac injury following AKI. We have demonstrated that AKI induced inflammation, mitochondrial dysfunction and pro-fibrotic factors in the heart (Figs. 2,3). In our study, proteomics analysis illustrated that SerpinA3N may be a potential manager of the ECM composition in the heart. Increased expression and activity of SerpinA3N, inhibits serine proteases thus impeding the clearance of extra ECM deposition (Fig. 4). These molecular consequences would negatively affect cardiac remodeling. We focused on a serine protease, Granzyme B (GZMB), which is a target of SerpinA3N, with the hypothesis that increased SerpinA3N protein expression inhibits the activity of GZMB and affects ECM composition maintenance in the heart tissue after AKI. We found that the GZMB activity was negatively correlated with SerpinA3N expression in the hearts of AKI as compared to Ctrl (Fig. 5).

Lastly, to show a direct SerpinA3N mediated GZMB activity inhibition, we employed H9c2 cells (Cardiomyoblasts) and induced SerpinA3N overexpression in these cells by treating these cells with H_2_O_2_ and assessed for secreted GZMB activity in the conditioned media from H9c2 cells cultures with and without a SerpinA3N inhibitor XAV939. Our results confirmed that while inhibiting SerpinA3N with XAV939, GZMB activity was increased (Fig. 6). Thus, confirming a direct SerpinA3N mediated inhibition of GZMB activity, which is otherwise difficult to accomplish in an *in vivo* setting given the XAV939 broad effect on other cells in the heart tissue.

These data suggest that AKI promotes inflammation, TGF-β1 mediated overexpression of SerpinA3N and pro-fibrotic factors, inflammatory cells mediated GZMB overexpression in the heart. Further, our results suggest that SerpinA3N is overexpressed and is activated in the heart of AKI mice. It inhibits the serine protease GZMB activity, which might lead to adverse cardiac remodeling as the balance between GZMB activity and ECM deposition is required for the critical maintenance of ECM and allow proper cardiac remodeling after acute cardiac injury due to AKI. Further studies using conditional SerpinA3N knockout mice in the heart tissue is required to confirm these findings and support the hypothesis that a decreased SerpinA3N expression in the heart allows for better cardiac remodeling after kidney injury.

## Author contributions

G.S.P. and R.L. designed experiments presented in the manuscript. G.S.P, R.N, N.Z. and K.W. performed experiments and analyzed the data presented in the manuscript.

N.H. and R.N. performed the mice IR-AKI surgeries. G.S.P wrote the manuscript. R.L, N.Z, N.S.B, G.V.H, L.L and J.B reviewed and edited the manuscript. All authors read and approved this manuscript for submission.

